# Peripheral nerve resident macrophages are microglia-like cells with tissue-specific programming

**DOI:** 10.1101/2019.12.19.883546

**Authors:** Peter L Wang, Aldrin KY Yim, Kiwook Kim, Denis Avey, Rafael S. Czepielewski, Marco Colonna, Jeffrey Milbrandt, Gwendalyn J Randolph, The Immunological Genome Project

**Author notes:** These authors contributed equally (Peter L Wang, Aldrin KY Yim).

## Abstract

Whereas microglia are recognized as fundamental players in central nervous system (CNS) development and function, much less is known about macrophages of the peripheral nervous system (PNS). Here we show that self-maintaining PNS macrophages share unique features with CNS microglia. By comparing gene expression across neural and conventional tissue-resident macrophages, we identified transcripts that were shared among neural resident macrophages as well as selectively enriched in PNS macrophages. Remarkably, PNS macrophages constitutively expressed genes previously identified to be upregulated by activated microglia during aging or neurodegeneration. Several microglial activation-associated and PNS macrophage-enriched genes were also expressed in spinal cord microglia at steady state. While PNS macrophages arose from both embryonic and hematopoietic precursors, their expression of activation-associated genes did not differ by ontogeny. Collectively, these data uncover shared and unique features between neural resident macrophages and emphasize the role of nerve environment for shaping PNS macrophage identity.

## Introduction

The significant role of resident neural macrophages in neuroinflammation and disease progression is increasingly appreciated in mouse models and individuals with neurodegeneration (1, 2, 3). Such advances, which largely rely on the interpretation of data from transcriptional analyses and human genome-wide association studies (GWAS) of Alzheimer’s Disease (AD) and other neurodegenerative conditions, have led to critical findings about cellular and molecular processes underlying such diseases (4, 5, 6). Most of these studies, however, have focused on resident macrophages in the brain (microglia) and, to a lesser extent, the spinal cord. Meanwhile, the transcriptional identity and functions of resident macrophages in the PNS remain mostly unknown.

The PNS consists of a multitude of neuronal networks that relay motor and sensory information between the CNS and the rest of the body (7). Though it has the capacity to regenerate, the PNS is also prone to injury and degeneration (8). Studies of PNS injury have shown that PNS macrophages play important roles for debris clearance, pain development, and regeneration (9, 10, 11). While the contribution of recruited monocytes cannot be excluded, these studies demonstrate the importance of PNS macrophages in nerve injury. Understanding the roles of these cells in homeostasis and disease may be broadly beneficial for resolving neuroinflammation.

In addition to monocyte-derived macrophages in nerve injury, there are also resident macrophages in the PNS at steady state (12, 13). While their residence in neuronal tissues is inherently microglia-like, PNS macrophages exist within a unique peripheral nerve microenvironment. Moreover, though it is known that CNS microglia are derived from the yolk sac during embryogenesis and depend on IL-34 for development (14, 15), the ontogeny of PNS macrophages remains unclear. Considering the growing interest in how tissue environment and ontogeny contribute to microglial identity and function in neurological diseases, we investigated these questions in peripheral nerve resident macrophages.

## Results

### Resident macrophages of the PNS

To examine resident macrophages in the PNS, we imaged a variety of nerve types at steady state using CX3CR1^GFP/+^ reporter mice. In these mice, GFP effectively labels microglia and has been shown to label nerve-associated macrophages in adipose and enteric tissues (16, 17). CX3CR1-GFP^+^ cells were found in dorsal root ganglia (DRG), vagal nerves (VN), subcutaneous fascial nerves (FN) and sciatic nerves (SN) (Fig. 1a). CX3CR1-GFP+ cells were located in the endoneurium and expressed colony-stimulating factor 1 receptor (CSF1R), also known as CD115 (Fig. 1b-c). Using flow cytometry, we found that CX3CR1-GFP^+^ cells also expressed the common macrophage marker CD64 (FcγR1) (18), as well as intermediate levels of CD45 (Fig. 1d-f). Thus, CX3CR1-GFP^+^ cells in peripheral nerves are indeed macrophages with resemblance to CNS microglia based on both endoneurial localization and surface marker expression.

**Figure 1.**
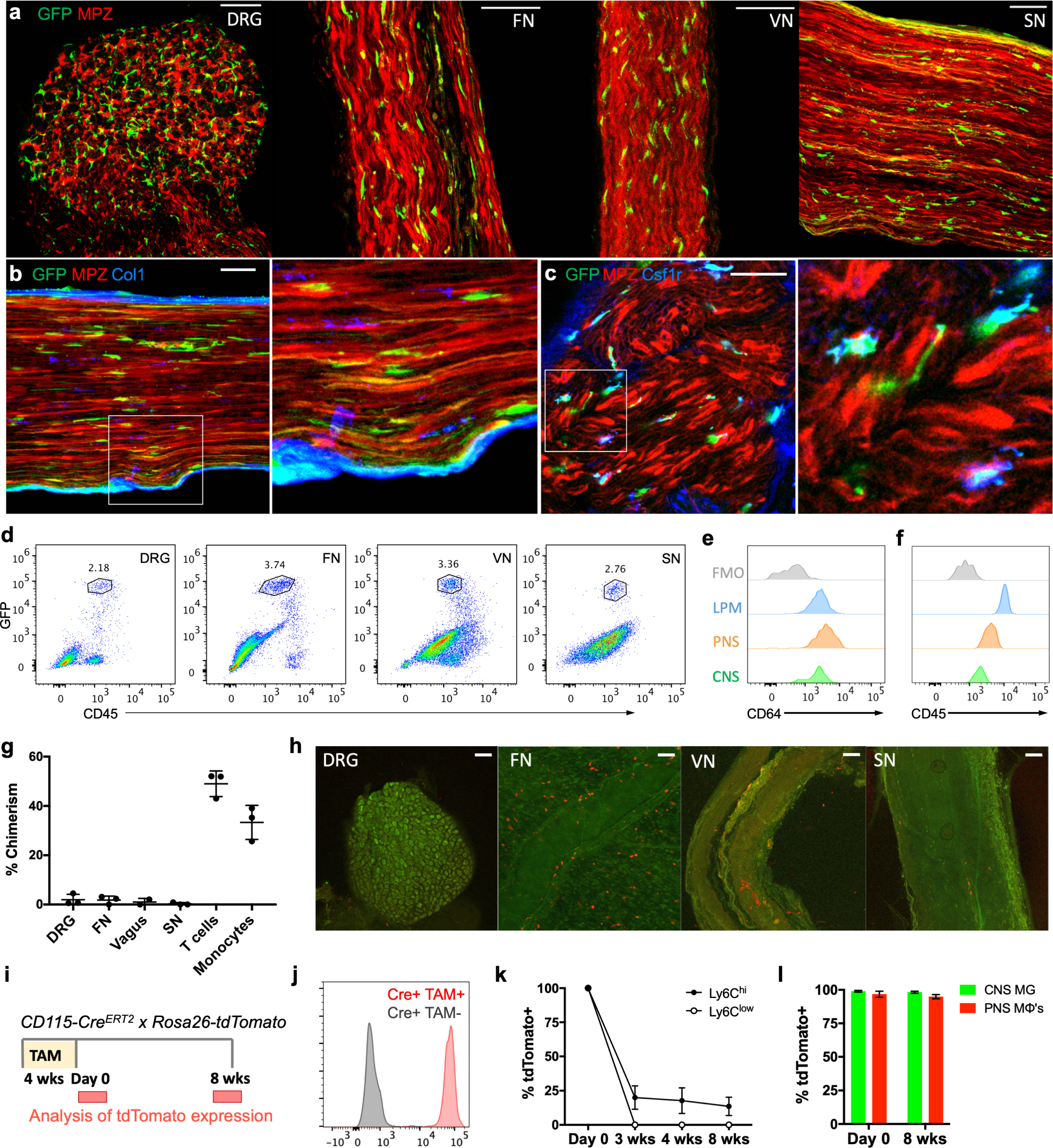
Identification and characterization of PNS resident macrophages. (a, b, c) Representative confocal imaging of peripheral nerves from CX3CR1^GFP/+^MPZ^tdTomato^ mice. (a) Images of whole mount dorsal root ganglia, subcutaneous fascia, vagal and sciatic nerves isolated from CX3CR1^GFP/+^MPZ^tdTomato^ mice. Scale bars are 50 microns (μm). (b) Endoneurial localization of CX3CR1^GFP/+^ cells in longitudinal sections of sciatic nerves; (c) Csf1r (blue) and CX3CR1^GFP^ (green) colocalization in sciatic nerve cross sections. Scale bars are 50 μm. (d) Flow cytometric gating of CX3CR1^GFP/+^ cells from peripheral nerve tissues. (e, f) Representative expression of CD64 and CD45 in CX3CR1^GFP/+^ cells compared to brain microglia, large peritoneal macrophages (LPMs), and fluorescence minus one (FMO) control. (g) Flow cytometric quantification of CD45.1 and CD45.2 chimerism in blood (total T cells or total monocytes) and nerves from 3 pairs of WT (CD45.1) and Lyz2Cre tdTomato (CD45.2) parabionts 10 weeks after joining; (h) Representative imaging in peripheral nerves from WT parabiont; Scale bars are 100 μm; (i-l) Analysis of tdtomato expression in tamoxifen-pulsed CSF1R^Mer-iCre-Mer^ x Rosa26-tdTomato mice. (i) Tamoxifen delivery schematic for fate mapping. Mice were given tamoxifen diet for 4 weeks and analyses for PNS macrophages and CNS microglia were performed at 0 days and 8 weeks after tamoxifen diet removal. (j) tdTomato expression by genotype from combined peripheral nerves in tamoxifen-fed mice. (k) Flow cytometric quantification of Ly6c high and Ly6C low monocytes from mice bled at 0 days, 3 weeks, 4 weeks, and 8 weeks after tamoxifen removal. (l) Flow cytometric quantification of CNS microglia (brain and spinal cord) and PNS macrophages (DRG, fascia nerve, vagus nerve, sciatic nerve) 0 days and 8 weeks following tamoxifen removal. (n=3 mice per time point) Data are mean +/− SEM.

To determine whether PNS macrophages depend on circulating precursors or are maintained via local signals, we performed parabiosis in CD45.1+ wild type and CD45.2+ Lyz2Cre x tdTomato^fl/fl^ mice and assessed the extent to which cells circulating from the parabiotic partner gave rise to PNS macrophages. Ten weeks after joining the parabionts, we found minimal exchange of PNS macrophages in all of the nerve types examined, while blood T cells and monocytes exchanged robustly (Fig. 1g; Supplementary Fig. 1). Indeed, most of the tdTomato+ cells that could be seen in the wild type parabiont were localized to the tissue surrounding the nerves (Fig. 1h). We also performed pulse chase labeling of PNS macrophages using tamoxifen-inducible CSF1R^Mer-iCre-Mer^ x tdTomato^fl/fl^ mice. In these mice, tdTomato expression persists in self-maintaining cells, but not in monocytes, which mostly turn over by 3-4 weeks after tamoxifen removal (19, 20). Heterozygous mice were fed tamoxifen diet for 4 weeks and then switched to normal diet (Fig. 1i). Just following tamoxifen removal, 96% of PNS macrophages, 99% of CNS microglia, and 100% of blood monocytes were tdTomato+ (Fig. 1j-l, Supplementary Fig. 2). Whereas only 20% of nonclassical and classical monocytes were still tdTomato+ by 3 weeks after tamoxifen removal, 98% of CNS microglia and 95% of PNS macrophages remained labeled up to 8 weeks following tamoxifen removal (Fig. 1l, Supplementary Fig. 2). Taken together, these results indicate that PNS macrophages are mostly self-maintained in adult mice.

### Transcriptional characterization of PNS macrophages

As we and others have previously demonstrated, unique gene expression profiles can be obtained in tissue-resident macrophage populations across tissue types (18, 21). To identify signature genes in peripheral nerve macrophages, we performed bulk RNA-seq to compare purified PNS resident macrophages sorted from the dorsal root ganglia, vagal, subcutaneous fascial, and sciatic nerves (Supplementary Fig. 3) with CNS microglia from the brain and spinal cord as well as previously characterized “conventional” macrophage populations from spleen, peritoneal cavity, and lungs. Global transcriptomic analysis revealed similarities within resident neural macrophages from both PNS and CNS, with PNS macrophages clustering more closely to CNS microglia than to conventional macrophages (Fig. 2a, Supplementary table 1). A substantial number of genes were uniquely enriched in PNS macrophages and CNS microglia compared to the other tissue-resident macrophages, including microglial signature genes *Tmem119*, *P2ry12*, *Siglech*, *Trem2*, and *Olfml3* (Fig. 2b, c). PNS macrophage-specific genes were also identified (Fig. 2b).

**Figure 2.**
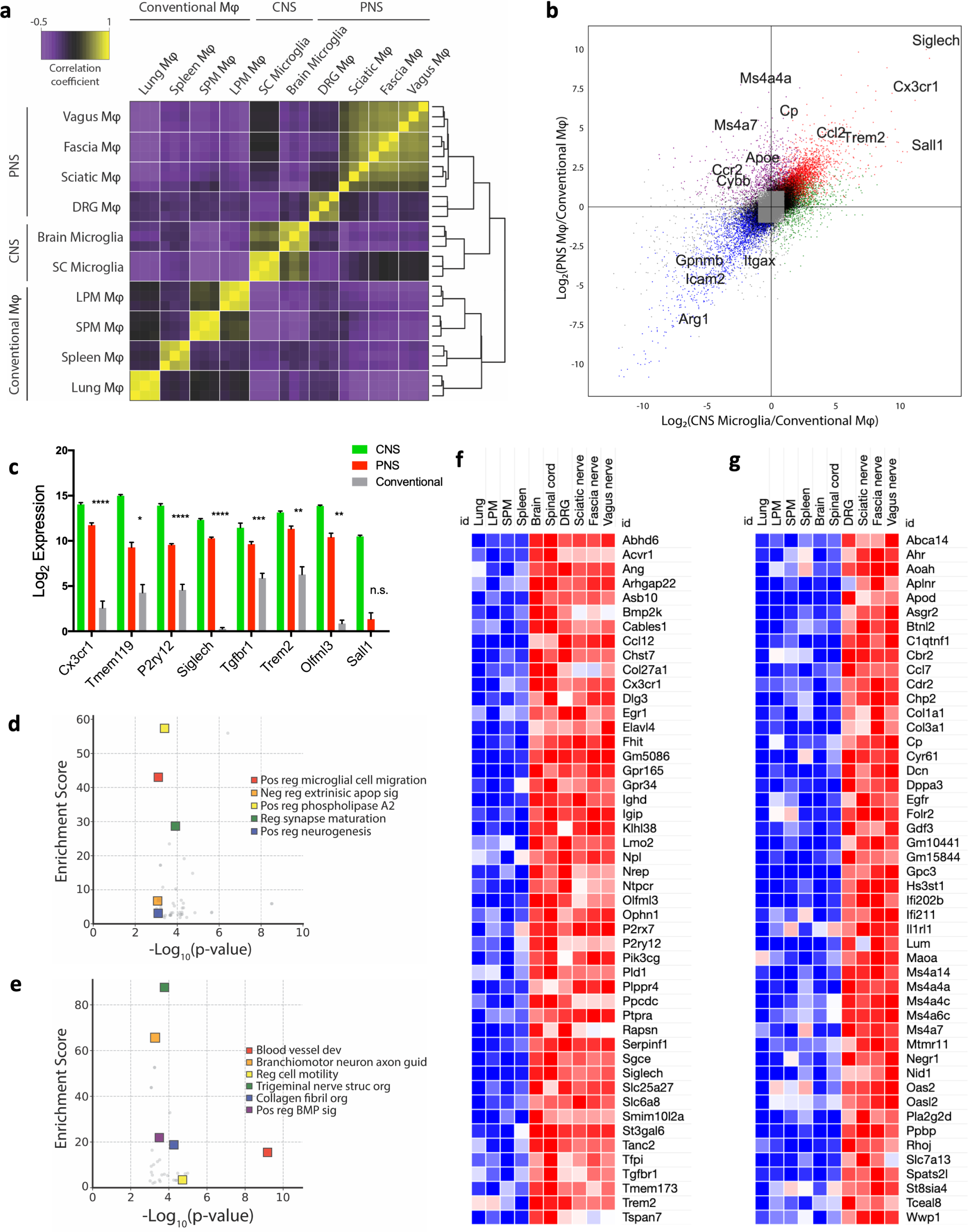
PNS macrophages express microglial transcripts as well as a unique signature. (a) Sample correlation plot showing global transcriptomic analysis and hierarchical clustering of resident macrophages from PNS, CNS, and conventional macrophages. Each box represents one replicate. 3 replicates comprised of up to 20 mice per replicate were included for each population. (b) Visualization of PNS macrophage unique transcripts (upper quadrant), CNS microglia unique transcripts (right quadrant), shared transcripts between PNS macrophages and CNS microglia (diagonal, top right quadrant), and conventional macrophages (bottom right quadrant). (c) Expression of microglial core transcripts in PNS macrophages compared with conventional macrophages. Multiple t tests. Data are mean +/− SEM. * P < 0.05, ** P < 0.01, *** P < 0.001, **** P < 0.0001. (d) GO analysis of genes enriched in PNS macrophages and CNS microglia. (e) GO analysis of genes enriched in PNS macrophages. (f) Transcripts expressed at least 4-fold higher in PNS macrophages and CNS microglia than conventional tissue-resident macrophages (*p* ≤ 0.05). (g) Transcripts expressed at least 4-fold higher in PNS macrophages than CNS microglia or conventional macrophages (*p* ≤ 0.05). (f, g) Each box represents average of three replicates.

To determine potential functions of PNS macrophages associated with their shared and unique gene expression profiles, we performed gene ontology (GO) analysis on transcripts that were common between PNS macrophages and CNS microglia and those that were specific to PNS macrophages (Supplementary table 2). Consistent with the idea that PNS macrophages may share functions with CNS microglia, pathway analysis identified functions including synaptic plasticity, microglial motility, and positive regulation of neurogenesis (Fig. 2d). Pathways that were unique to PNS macrophages included angiogenesis, collagen fibril organization, regulation of BMP signaling, and peripheral nerve structural organization and axon guidance (Fig. 2e).

Next, we identified transcripts that were 4-fold or more enriched in CNS microglia and PNS macrophages relative to their expression in all other conventional macrophage populations (Fig. 2f). These upregulated genes included *Abhd6*, *Ophn1*, *P2rx7*, *Pld1*, *Sgce*, *Tgfbr1, Tfpi*, and *Tmem173*. We also identified several genes that were downregulated in CNS microglia and PNS macrophages (Supplementary Fig. 4). Notably, the transcriptional regulator that defines microglial identity and function, *Sall1*, was not expressed by PNS macrophages (Fig. 2c). Modest detection of *Sall1* in the DRG could not be corroborated by further analysis (Supplementary Fig. 5). This may reflect unique adaptations in PNS and CNS macrophages. Indeed, we identified 72 genes, including *Sall1*, that were highly specific to CNS microglia (Supplementary Fig. 6).

To examine unique gene expression in PNS macrophages, we identified transcripts that were at least 4-fold higher or lower in PNS macrophages compared to other resident macrophages, including CNS microglia (Fig. 2g, Supplementary Fig. 7). We found 48 genes specifically enriched in PNS macrophages, including *Aplnr*, *Cp*, *Il1rl1*, *Maoa*, *Pla2g2d*, and *St8sia4*, as well as interferon-induced genes *Ifi202b*, *Ifi211*, and *Oas2*. We also identified *Ms4a14*, *Ms4a4a*, *Ms4a4c*, *Ms4a6c*, and *Ms4a7*. These signatures reveal unique transcriptional programming in PNS macrophages and may provide clues about their involvement in neuronal health and disease.

We next investigated transcriptional differences within PNS macrophage populations. Using a 4-fold cutoff, we identified 24 genes enriched in sciatic nerve macrophages, 23 genes enriched in fascial nerve macrophages, and 12 genes enriched in vagal nerve macrophages (Supplementary Fig. 8a). We observed similar numbers of downregulated genes in each population (Supplementary Fig. 8b). We also compared nerve-resident macrophages to those residing within the dorsal root ganglion and found that they were significantly different, with many differentially expressed transcripts. Therefore, we re-analyzed this data using a more stringent 8-fold cutoff and identified 79 upregulated genes and 52 downregulated genes (Supplementary Fig. 8c, d). These results suggest that while peripheral nerve macrophages are transcriptionally similar, significant differences exist between those adjacent to axons and those residing close to neuronal cell bodies.

### PNS macrophages express microglial activation genes

To investigate differentially expressed genes (DEGs) within resident neural macrophages, we refined our analysis to CNS microglia and PNS macrophages (Fig. 3a). We identified 396 genes enriched in PNS macrophages and 180 genes enriched in CNS microglia (Supplementary Table 3). Since the upregulation of MS4A family and interferon-induced genes has been reported to characterize aged and neurodegenerative disease-associated microglia (6, 22, 23), we wondered if PNS macrophages expressed other genes associated with microglial activation. By cross-referencing published data, we determined the number of connections between disease-associated genes that were upregulated in activated microglia from aging, phagocytic, and neurodegenerative conditions and neural macrophage-enriched genes from either PNS macrophages or CNS microglia (Fig. 3b). We found 148 disease-associated genes that were enriched in PNS macrophages compared to 17 that were enriched in CNS microglia (Fig. 3b, Supplementary table 4). From the highest connectivity groups (6–4), we identified 25 genes that were significantly higher in PNS macrophages, including *Ch25h*, *Anxa2*, *Cd52*, *Ifitm3*, *Cybb*, *Fxyd5*, *Igf1*, and *Apoe* (Fig. 3c).

**Figure 3.**
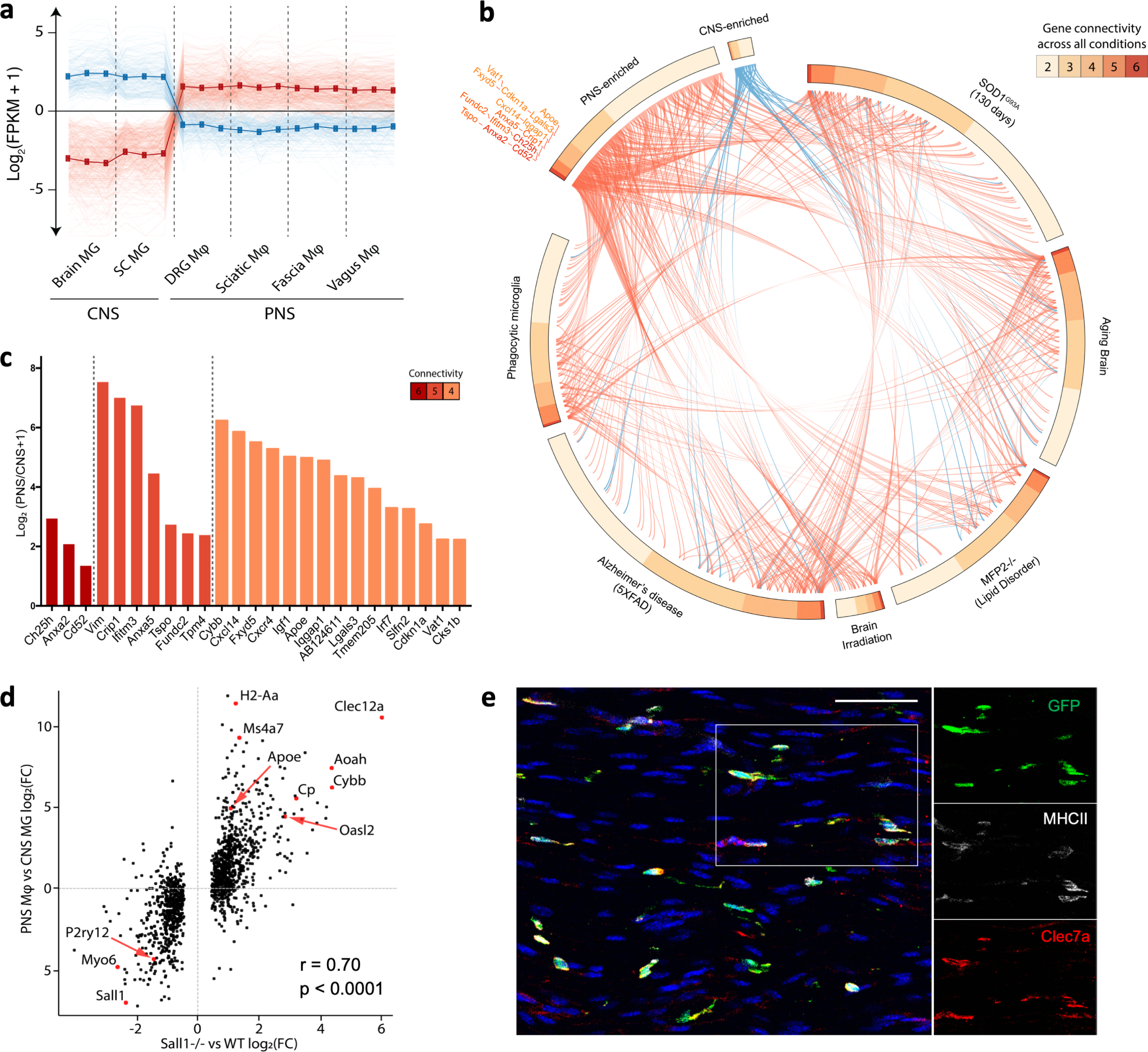
PNS macrophages constitutively express transcripts associated with activated microglia. (a) Expression pattern of DEGs defined as PNS-enriched (red) or CNS-enriched (blue). CNS microglia includes brain and spinal cord and PNS macrophages include DRG, vagal, fascial, and sciatic nerves. (b) Circos plot showing the number of connections between GSEA-scored genes from microglia in 6 neurodegenerative and aging-associated conditions as defined in Krasemann et al. and neural macrophage-enriched genes from either PNS macrophages or CNS microglia (gene connectivity). (c) PNS-enriched genes from connectivity groups 6-4 expressed as Log_2_ fold change (PNS macrophages/CNS microglia). (d) Expression plot comparing PNS macrophage-enriched genes (expressed as PNS macrophage/CNS microglia Log2FC) and Sall1^−/−^ microglia-enriched genes (expressed as KO/WT Log2FC) from Buttgereit et al. r, correlation coefficient; p, p-value for linear regression analysis. (e) Representative immunohistochemistry in sciatic nerves of CX3CR1^GFP/+^ mice showing DAPI (blue), GFP (green), MHCII (white), and Clec7a (red).

Microglia lacking certain genes for homeostatic regulation have also been found to shift their gene expression towards an activated phenotype (24, 25). Sall1 has been identified as a transcriptional regulator of microglia identity and function, with Sall1^-/-^ microglia resembling inflammatory phagocytes (24). Since PNS macrophages did not express Sall1 at steady state, we examined whether genes that are reportedly dysregulated in Sall1^-/-^ microglia showed the same pattern of expression in PNS macrophages. Indeed, we found a high correlation between genes enriched in PNS macrophages and Sall1^-/-^ microglia, including *Apoe*, *H2-Aa*, *Ms4a7*, *Clec12a*, *Aoah*, and *Cybb* (Fig. 3c). These data suggest that PNS macrophages may share common genetic regulators with CNS microglia.

Since microglia activation may occur under cell sorting conditions (26), we were concerned that the activation signature in PNS macrophages might be attributed to tissue preparation. Thus, we stained freshly fixed peripheral nerves for Clec7a and MHCII, which are induced in microglia across many activation states (6, 25, 27). Resting PNS macrophages were clearly marked by Clec7a and MHCII (Fig. 3e), suggesting that the signature obtained in PNS macrophages is not a technical artifact. Taken together, these data show that PNS macrophages constitutively express a wide array of microglial activation genes and imply a shared yet microenvironment-sensitive regulation of gene expression in resident neural macrophages.

### PNS to CNS zonation in resident neural macrophages

Given the difference in gene expression between PNS macrophages and CNS microglia, we wanted to further discriminate the influence of microenvironment on neural macrophage identity. Thus, we examined spinal cord microglia, which reside in a distinct microenvironment from brain microglia. Interestingly, we found a set of genes that were high in PNS macrophages, intermediate to high in spinal cord microglia, and low in brain microglia (PNS to CNS zonation) (Fig. 4a, Suppl. Table). These included PNS macrophage-specific genes (*Cp*, *Il1rl1*, *Maoa*, and *Cdr2*), microglial activation genes (*Clec7a*, *Spp1*, *Lpl*, *Axl*, *Ms4a4c*, and *Ms4a6c*), interferon-induced genes (*Ifi204*, *Ifi207*, *Ifi209*, and *Oasl2*), mitochondrially encoded genes (*mt-Nd1*, *mt-Nd2*, *mt-Nd4*, *mt-Nd5*, and *mt-Nd6*), and several transcription factors (*Hivep2*, *Zfp704*, and *Rbpj*) (Fig. 4b, c). We confirmed Clec7a expression in spinal cord microglia by immunostaining (Fig. 4c). Importantly, microglia from spinal cord and brain did not significantly differ by expression of homeostatic genes Sall1, Olfml3, and Tmem119 (Supplementary Fig. 9). In fact, certain microglial genes, including Tgfbr1 and P2ry12 were more highly expressed in spinal cord compared to brain microglia. These findings suggest that transcriptional programs underlying PNS macrophages and activated microglia may be present during normal physiological conditions and further support the role of nerve environment for specifying neural macrophage identity.

**Figure 4.**
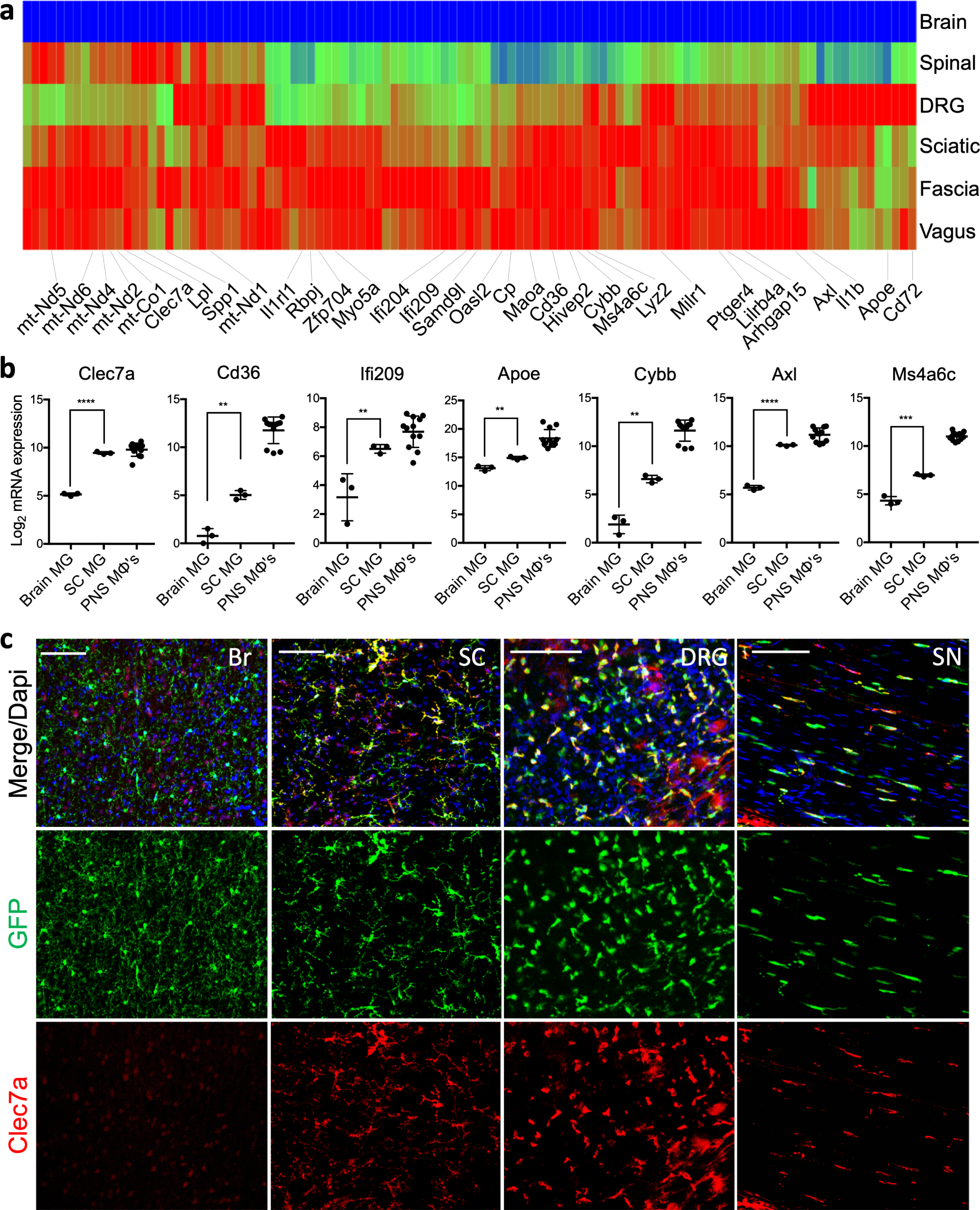
Zonation of PNS macrophage-enriched transcripts in neural resident macrophages. (a) Heat map showing genes corresponding to PNS to CNS zonation pattern (PNS MΦ’s high, SC MG high/intermediate, brain MG low). Each box represents average from 3 replicates. (b) Gene expression analysis of individual genes following PNS to CNS zonation pattern. Each dot represents one replicate. Unpaired t test. Data are mean +/− SD. * P < 0.05, ** P < 0.01, *** P < 0.001, **** P < 0.0001, (c) Representative Clec7a staining in brain (Br), spinal cord (SC), dorsal root ganglia (DRG), and sciatic nerve (SN) of CX3CR1^GFP/+^ mice. Scale bars are 100 μm.

### Ontogeny of PNS resident macrophages

In addition to microenvironmental cues, ontogeny may also play an important role for specifying microglial identity and function. Specifically, it has been shown that, compared to naturally occurring microglia with yolk sac origin, monocyte- and hematopoietic stem cell (HSC)-derived microglia display a more activated signature (28, 29). Thus, we sought to determine PNS macrophage ontogeny by examining Flt3Cre x LSL-YFP^fl/fl^ reporter mice (Fig. 5a), a strain that labels fetal monocyte and adult HSC-derived hematopoietic multipotent progenitors and their progeny (30, 31). We found that approximately 26% of PNS macrophages across nerve types appeared to be embryonically derived (YFP-) and 74% of PNS macrophages were HSC-derived (YFP+) (Fig. 5b, c).

**Figure 5.**
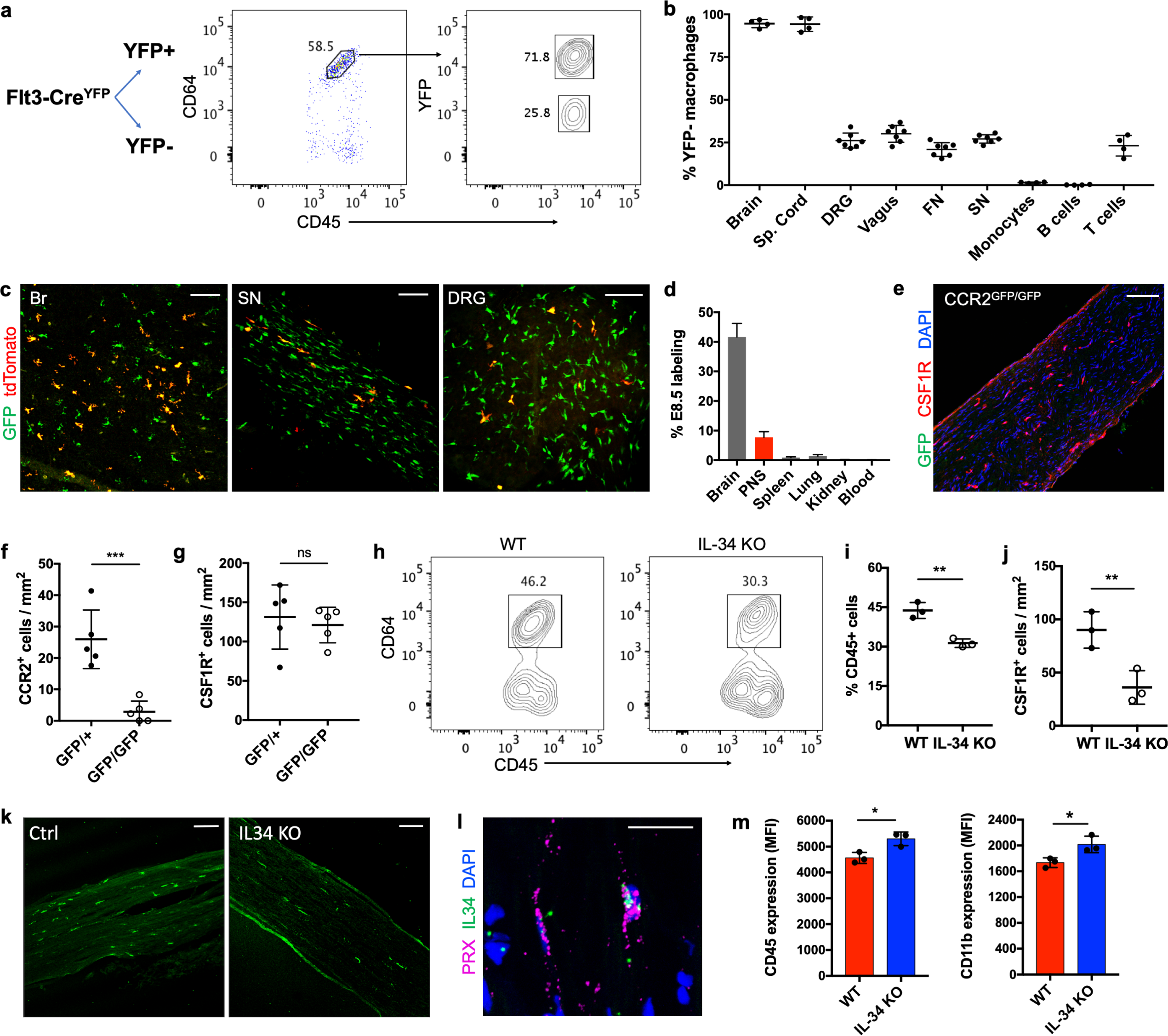
Shared and distinct developmental programs in PNS macrophages. (a) Schematic and flow cytometric gating for analysis of YFP expression in PNS macrophages from Flt3-Cre LSL-YFP sciatic nerves. (b) Comparison of YFP-macrophage ratios (expressed as % YFP-cells of population) across neural resident macrophage populations, monocytes, B cells, and T cells. (*n* = 4-7 mice per group) (c) Image showing E8.5 tdTomato labeling and GFP expression in brain (left panel), sciatic nerve (middle panel), and DRG (right panel) of CSF1R^Mer-iCre-Mer^ x tdTomato^fl/fl^ x CX3CR1^GFP/+^ mice. Scale bar, 100 μm. (d) Quantification of E8.5 tdTomato labeling in brain, PNS, spleen, lungs, kidney, and blood from newborn pups. Several images were taken and analyzed from each tissue and averages from each group were plotted. (n=2 mice for blood analysis, n=3 mice for tissue analysis). (e) Histological representation of PNS macrophages in CCR2^GFP/GFP^ sciatic nerves. Scale bar, 100 μm (f) Quantification of CCR2+ CSF1R+ macrophages in sciatic nerves in CCR2^GFP/+^ and CCR2^GFP/GFP^ mice (n=5 mice per group). (g) Quantification of total CSF1R+ macrophages in CCR2^GFP/+^ and CCR2^GFP/GFP^ mice (n=5 mice per group). (h) Flow cytometric gating and (i) analysis of sciatic nerve macrophages from WT and IL-34 KO mice. (n=3 mice per group). (j) Histological quantification of sciatic nerve macrophages in WT and IL-34 KO mice. At least 3 images were analyzed and averaged for each nerve (*n* = 3 mice per group). Scale bar, 100 μm (k) Representative imaging of WT and IL-34 KO sciatic nerves. CSF1R staining in green. (l) In situ hybridization in WT sciatic nerve showing IL-34 localization with Prx-expressing Schwann cell. Scale bar, 50 μm (m) Flow cytometric representation of CD11b and CD45 expression changes in PNS macrophages from WT (red) and IL-34^LacZ/LacZ^ (blue) mice. Data is reflective of all replicates examined (n=3 mice per group). Each dot represents one mouse. All data are analyzed by unpaired t test. Data are mean +/− SD. * P < 0.05, ** P < 0.01, *** P < 0.001, **** P < 0.0001; ns, not significant.

To further examine the contribution of embryonic precursors to PNS macrophages, we performed fate mapping using CSF1R^Mer-iCre-Mer^ x tdTomato^fl/fl^ x CX3CR1^GFP/+^ double reporter mice, which allows simultaneous visualization of tamoxifen-pulsed CSF1R-expressing macrophages and resident Cx3cr1+ macrophages. After giving tamoxifen at embryonic day 8.5 (E8.5), a developmental time point that labels yolk-sac derived macrophages, we checked newborn pups for labeling. The PNS was fully populated with CX3CR1-GFP/+ macrophages at birth. In line with previous observations (32), we observed partial labeling in 42% of brain microglia (Fig. 5c). Labeling in PNS macrophages was 8% from sciatic nerve and DRG, or roughly one-fifth of the entire PNS macrophage population when normalized to microglia (Fig. 5c). Less than 1% of macrophages were tdTomato+ in spleen, kidney, and blood, and only 3% of lung macrophages were tdTomato+ (Fig. 5c, Supplementary Fig. 12). These results demonstrate that the PNS is fully populated by macrophages at the time of birth and that a subset of these cells are derived from yolk sac progenitors.

While monocyte entry into injured nerves depends on Ccr2 (33, 34), it is not known whether Ccr2 is required for seeding or maintaining the PNS macrophage niche. To address this question, we quantified CSF1R+ macrophages in sciatic nerves of Ccr2 knock-in (Ccr2^GFP/GFP^) and control (Ccr2^GFP/+^) mice. We observed no difference in PNS macrophage numbers between Ccr2 knockouts and controls (Fig 5e-g). However, while GFP^+^ cells were present in sciatic nerves from Ccr2^GFP/+^ mice, GFP^+^ cells were totally absent from Ccr2^GFP/GFP^ mice (Fig. 5e, f). These observations suggest that Ccr2-dependent monocyte entry is not required to fill or maintain the PNS macrophage niche, but may contribute to a modest subset of PNS macrophages at steady state.

Given the resemblance between microglia and PNS macrophages, we wondered if IL-34, an alternative ligand for CSF1R that contributes to CNS microglia development (15), is likewise important for PNS macrophage development. We thus quantified PNS macrophages in sciatic nerves of IL-34 deficient mice (IL-34^LacZ/LacZ^) and found a 30% reduction in PNS macrophage percentage by flow cytometry (Fig. 5h-i) and nearly 60% reduction in total numbers in IL-34 KO mice when quantified in sections (Fig. 5j-k), underscoring significant dependence on IL-34. To determine what cells in the nerve were producing IL-34, we checked IL-34 transcriptional localization by in situ hybridization. Surprisingly, we found that Prx-expressing myelinating Schwann cells were a source of IL-34 (Fig. 5l). In the absence of IL-34, PNS macrophages showed increased surface marker expression of CD45 and CD11b, suggesting a shift in phenotype away from microglial characteristics (Fig. 5m). Taken together, these results reveal shared and unique developmental programs between PNS macrophages and CNS microglia.

### Nerve environment shapes PNS macrophage signature

To investigate the extent to which PNS macrophage identity is specified by ontogeny or nerve environment, we individually sorted YFP-embryonic-derived and YFP+ HSC-derived PNS macrophages from sciatic nerves of Flt3Cre LSL-YFP^fl/fl^ mice and performed single cell RNA-seq (Fig. 6a). We captured a total of 935 YFP-cells and 3,186 YFP+ cells. Unsupervised clustering analysis of all 4,121 cells revealed 5 separate clusters, with the majority of PNS macrophages belonging to one major group (clusters 1, 2, and 3) and the remainder falling into two smaller clusters (4 and 5) (Fig. 6b, Supplementary table 5).

**Figure 6.**
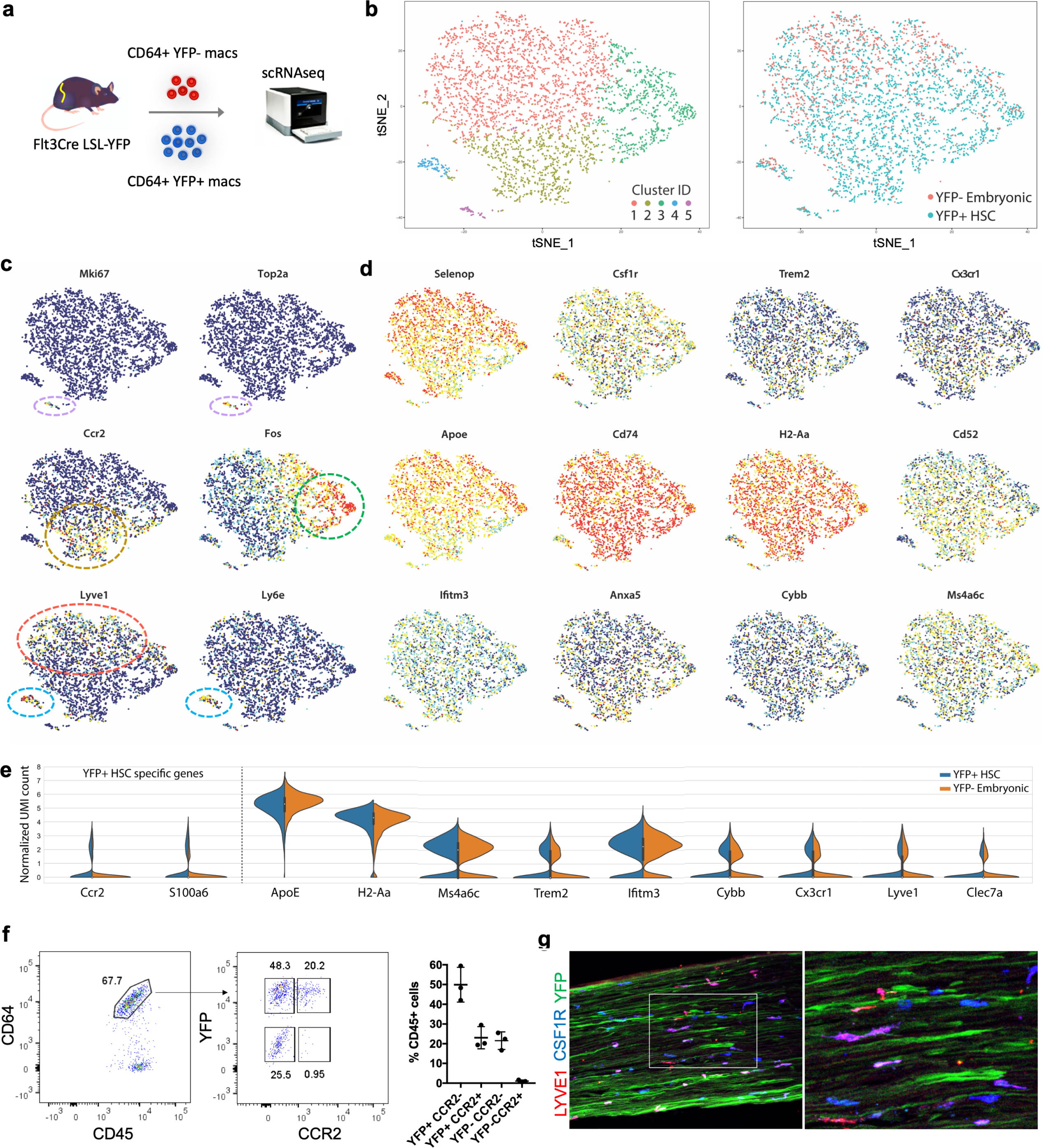
Nerve environment shapes transcriptional identity of PNS macrophages. (a) Schematic for isolation and separate single cell RNA sequencing of YFP+ HSC-derived and YFP-embryonic macrophages from sciatic nerves of Flt3-Cre LSL-YFP^fl/fl^ mice. Two single cell libraries from YFP+ and YFP-macrophages were prepared using the 10X single cell RNA-seq platform (b) t-SNE plot of 4,121 CD64+ CD45int cells from pooled sciatic nerves (n=18) showing unsupervised clustering (left panel) and overlay of YFP+ (blue) and YFP-(red) populations (right panel). (c) t-SNE plots depicting distribution of transcripts across combined YFP+ HSC-derived and YFP-embryonic PNS macrophages. (d) Violin plots of marker gene expression in YFP+ HSC-derived and YFP-embryonic groups. (e) Flow cytometric identification of CCR2+ macrophages in a subset of YFP+ macrophages. Gating is representative of at least 7 nerves examined over three separate experiments (f) Representative imaging of a subset of Lyve1+ macrophages in sciatic nerve.

While we observed a significant overlap of YFP- and YFP+ macrophages in clusters 1, 3, 4, and 5, cluster 2 contained mostly YFP+ macrophages (Fig 6b). Interestingly, cluster 2 was defined by *Ccr2* (and *S100a6*) expression, which is consistent with this subset arising from circulating precursors (Fig. 6c). Indeed, we confirmed by flow cytometry that CCR2+ PNS macrophages were only found in the YFP+ fraction (Fig. 6f). We also observed varying heterogeneity between the overlapping clusters (Fig. 6c). For instance, cluster 5 was easily distinguished by proliferation genes *Mki67* and *Top2a*. Cluster 4 was also relatively distinct and showed enrichment for *Ly6e*, *Ninj1*, *Retnla*, and *Wfdc17*, potentially representing a previously undescribed activation state. Cluster 3 selectively expressed early activation genes *Fos*, *Jun*, and *Egr1* and may represent cells from clusters 1 and 2 undergoing acute activation from cell preparation and sorting. Cluster 1, the least distinct population, was slightly enriched for *Lyve1* expression compared to cluster 2 (Fig. 6c). We confirmed Lyve1 expression in a subset of PNS macrophages by immunostaining (Fig 6g).

Despite the identification of separate clusters in our data, we observed no obvious difference in the expression of PNS macrophage-enriched or microglial activation-associated transcripts between the main clusters. Specifically, *Apoe*, *Cp*, *H2-Aa*, *Cd74*, *Ms4a6c*, *Ifitm3*, *Anxa5*, *Cybb*, and *Cd52* were nearly identical between clusters 1, 2, and 3 (Fig. 6d). We also observed no difference in *Cx3cr1* and *Trem2* between these clusters. Importantly, all of these genes were equally expressed between YFP- and YFP+ macrophages (Fig. 6e). We conclude that embryonic- and HSC-derived PNS macrophages are transcriptionally similar and that the nerve environment confers a predominant effect over developmental origin on PNS macrophage identity.

## Discussion

PNS-resident endoneurial macrophages are a previously unprofiled population of neural resident macrophages. Here we characterized the transcriptomes of PNS resident macrophages across various nerve types spanning both axons and cell bodies and innervating a wide range of somatic targets. We show that self-maintaining PNS macrophages possess shared gene expression programs with CNS microglia as well as a unique set of genes shaped by the nerve environment. The convergence of both HSC-derived and embryonic PNS macrophages into a single population at steady state underscores the importance of tissue environment for specifying PNS macrophage identity.

Our findings demonstrate that PNS macrophages share a significant overlap in gene expression with CNS microglia. This resembles recent findings that show microglial genes being expressed in nerve-associated macrophages in gut and adipose tissues (16, 17). While PNS macrophages reside in the nerve proper, there are likely common tissue factors that induce nerve-imprinted signatures in macrophages. One such factor may be transforming growth factor beta (TGF-β). TGF-β signaling is vital for the expression of homeostatic genes in microglia (2, 3). The higher expression of *Tgfbr1* in PNS macrophages and CNS microglia at steady state supports the role of TGF-β signaling in neural resident macrophages.

We were surprised that *Sall1* expression was so low in PNS macrophages. This likely represents a key difference in PNS and CNS nerve environments. The identification of additional genes that were purely expressed in CNS microglia may provide clues for the basis of this difference. Future studies should address genetic and environmental factors that regulate *Sall1* expression and examine whether it is expressed in a context-dependent manner in PNS macrophages.

The constitutive expression of a broad set of microglial activation genes in PNS macrophages and, to a lesser degree, in spinal cord microglia suggests that such programs may be intrinsic to their functioning in distinct neuronal environments. Indeed, it was found that sympathetic nerve-associated (SAM) macrophages and macrophages at CNS interfaces also appear to be constitutively activated (16, 35, 36). Additionally, an overlap in many of the same activation genes are induced in microglia localized to white matter tracts during brain development (37). However, this is the first time that endoneurial macrophages in the PNS have been shown to possess activation signatures at steady state. Further studies are needed to determine whether immune response, phagocytic and tissue remodeling programs are more prevalent at steady state.

While the connection between neurodegeneration and immune response has become increasingly apparent, how distinct expression patterns relate to neuronal disease, development, and homeostasis remains a mystery. Ontogeny seems to be an important factor, as shown by several studies that found upregulation of disease-associated genes and neurotoxic functions in transplanted cells with hematopoietic origin (28, 29). Our results show that both embryonic and HSC-derived PNS macrophages exist as a transcriptionally similar population in adult nerves regardless of origin, thus suggesting a more important role for nerve environment in programming neural resident macrophages at specific sites. Moreover, since PNS macrophages are a naturally occurring population, they serve as a useful standard of comparison in the context of normal physiological function.

Up to now, the brain and spinal cord of the CNS are the only tissues thought to possess microglia. Our comparison of resident macrophages from both PNS and CNS challenges this notion and supports the idea that microglial features and transcriptional programs are shared in macrophages across the nervous system. The increasing amount of transcriptomic data on microglia and nerve-associated macrophages from different tissues and across organs presents opportunities for novel discoveries that may be applied to neurodegenerative conditions and beyond.

## Supplemental Figures

Supplemental figure 1) Analysis of resident macrophage chimerism in CD45.1 wild type and CD45.2 Lyz2-Cre tdTomato parabionts. Representative flow plots of CD45.1 and CD45.2 expression in nerve macrophages from CD45.2 parabiont.

Supplemental Figure 2) Flow cytometric analysis of blood and neural resident macrophages in pulse chase experiment (related to Figure 1j-l). Representative flow gating of CNS microglia, PNS macrophages, and monocytes at day 0 and 8 weeks after tamoxifen removal.

Supplemental figure 3) Gating strategy for identifying PNS macrophages (related to Figure 1) and for double-sorted populations used in bulk-RNA sequencing.

Supplemental figure 4) Uniquely downregulated genes in neural resident macrophages. Heat map of mRNA transcripts fourfold or more downregulated in CNS microglia and PNS macrophages compared to conventional tissue-resident macrophages.

Supplemental figure 5) Sall1 expression in PNS macrophages. Flow cytometry analysis of GFP expression in brain microglia, DRG macrophages, and sciatic nerve macrophages isolated from Sall1-GFP/+ mice (red) and wild type controls (blue).

Supplementary Figure 6) Identification of unique signatures in CNS microglia. Heat map and gene list reveal transcripts selectively enriched in brain and spinal cord microglia fourfold or more compared to PNS and conventional macrophages.

Supplementary Figure 7) Downregulated genes in PNS macrophages. Heat map of genes that are downregulated fourfold or more in PNS macrophages compared to all other populations combined.

Supplementary Figure 8) Unique gene expression patterns in individual PNS macrophage populations. Heat map of upregulated and downregulated mRNA transcripts in single PNS macrophage populations by fourfold or more in sciatic, fascial, and vagal nerves and eightfold or more in DRG relative to their expression in the remaining three populations combined.

Supplementary Figure 9) Expression of microglial homeostatic genes in spinal cord and brain microglia. Log_2_ expression microglial homeostatic genes in brain (green) and spinal cord (magenta) microglia. Multiple t tests. Data are mean +/− SEM with adjusted p-value shown.

Supplementary Figure 10) Flow cytometric gating of blood and CNS populations in Flt3-Cre LSL-YFP. Gating strategy for determining YFP-percentages in monocytes, B cells, T cells, brain microglia, and spinal cord microglia (related to Figure 5b).

Supplementary Figure 11) Embryonic 8.5 labeling in blood and non-neuronal tissues. Representative flow cytometric gating and imaging of tdTomato labeling in blood, spleen, lung, and kidney of CSF1R^Mer-iCre-Mer^ x tdTomato^fl/fl^ x CX3CR1-GFP/+ newborn pups pulsed with tamoxifen at E8.5. Scale bar, 100 μm.

Supplementary Figure 12) CCR2 is not required for seeding and maintaining PNS macrophages (related to Figure 5). Representative imaging of total CSF1R+ and CCR2+ macrophages in sciatic nerve sections of CCR2^GFP/+^ and CCR2^GFP/GFP^ mice. Scale bar, 100 μm.

Supplementary Figure 13) Schwann cells are a source of IL-34 (related to Figure 5l). In situ hybridization in sciatic nerve showing colocalization of IL-34 (green) and Prx (magenta). Scale bar, 50 μm.

Supplementary Figure 14) PNS macrophages in IL-34 KO undergo phenotypic shift in CD45 and CD11b expression (related to Figure 5m). Representative flow gating and expression (median fluorescence intensity, MFI) of CD45 and CD11b in sciatic nerve macrophages from control and IL-34 KO mice. (n=3 per group).

Supplementary Figure 15) Quantification of CCR2+ subsets in YFP+ and YFP-macrophages in Flt3-Cre LSL-YFP mice (related to Figure 6f). Representative gating scheme for identifying CCR2+ macrophages in Flt3-Cre LSL-YFP mice. (n = 3).

## Materials and Methods

### Experimental animals

Mouse care and experiments were performed in accordance with protocols approved by the Institutional Animal Care and Use Committee at Washington University in St. Louis under the protocols 20170154 and 20170030. Mice were kept on a 12-hour light dark cycle and received food and water *ad libitum*. The following strains were used: C57/B6 CD45.1 (stock no. 002014), CX3CR1^GFP/+^ (Cx3cr1tm1Litt/LittJ; stock no. 008451), LysM^cre/+^ (stock no. 004781), and Rosa^Lsl-Tomato^ (stock no. 007905) mice were purchased from Jackson Laboratory (JAX) and bred at Washington University. CSF1R^Mer-iCre-Mer^ mice (39) were obtained from JAX (stock no. 019098) and backcrossed fully to C57BL/6 background using the speed congenics core facility at Washington University, MPZ-Cre (40), Flt3-Cre LSL-YFP^fl/fl^ (31) mice were kindly provided by D. Denardo, CCR2^gfp/+^ (B6(C)-Ccr2^tm1.1Cln^/J, (JAX stock no. 027619) mice were kindly provided by K. Lavine, IL34^LacZ/LacZ^ mice (15) were kindly provided by M. Colonna, and Sall1-^GFP^ mice (24, 41) were kindly provided by M. Rauchman.

### IHC

For whole mount imaging, samples were harvested and immediately stored in 4% paraformaldehyde (PFA) containing 40% sucrose overnight. Samples were then washed in PBS, blocked, stained, and imaged. For frozen sections, samples were harvested, stored in PFA/sucrose, and embedded into OCT. 15 micron cuts were made for sciatic nerves. Sections were then blocked in 1% BSA, stained, and imaged. Antibodies to the following proteins were used: anti-GFP, Clec7a (Clone R1-8g7), CSF1R (R&D Systems, Accession # P09581), MHCII (IA/IE clone M5/114.15.2), LYVE1 (ab14917).

### Preparation of single-cell suspensions

For blood, mice were bled from the cheek immediately before sacrifice and cells were prepared as previously described. For nerves and all other tissues, mice were sacrificed and perfused with PBS. Nerves were harvested and kept on ice until dissociation. For ImmGen samples, nerves from 4-20 mice were pooled for each replicate. Cells were then incubated with gentle shaking for 20 minutes in digestion media containing collagenase IV, hyaluronidase, and DNAse. Cells were then washed and filtered through 70 μm cell strainers. For brain and spinal cord, myelin was removed using a 40/80% Percoll gradient.

### Flow cytometry

Single-cell suspensions were stained at 4°C. Dead cells were excluded by propidium idodide (PI). Antibodies to the following proteins were used: B220 (clone RA3-6B2), CCR2 (clone SA203G11), CD3e (clone 145-2C11), CD4 (clone RM4-5), CD8 (clone 53-6.7), CD11b (clone M1/70), CD16 (clone 2.4G2), CD45 (clone 30-F11), CD64 (clone X54-5/7.1), CD115 (clone AFS98), GR1 (clone 1A8), and Ly-6C (clone HK1.4). Cells were analyzed on a LSRII flow cytometer (Becton Dickinson) and analyzed with FlowJo software.

### Cell sorting

For bulk RNAseq, cells from 6-week-old male mice were double sorted on a FACSAria II (Becton Dickinson) for a final count of 1000 cells into lysis buffer according to the ImmGen Consortium standard operating protocol. Tissues were collected into culture medium on ice and subsequently digested with collagenase IV, hyaluronidase, and DNAse. Following digestion, samples were washed and kept on ice until sorting. The sort was repeated so that all cells were sorted twice, with a minimum of 1000 cells recovered in the second sort, sorting the second time into 5 µl TCL buffer containing 5% BME. Samples were kept at −80° C until further processing. For sorting in preparation of the Flt3-Cre LSL-YFP single cell Seq experiments, sciatic nerves from 19 male mice aged 10-12 weeks were combined and CD64+ CD45int macrophages were sorted individually into YFP+ and YFP-groups, yielding 18,000 and 5,000 cells, respectively. Both groups were immediately run on the 10x Genomics Chromium Controller according to the manufacturer’s protocol.

### Parabiosis

Parabiotic pairs were generated as previously described (38). C57/B6 (CD45.1) mice were paired with Lyz2Cre tdTomato (CD45.2) mice. Mice were injected with buprenorphine-SR subcutaneously prior to surgery. After 10 weeks, mice were sacrificed and nerve tissue was examined by flow cytometry and imaging to detect hematopoietic contribution to PNS resident macrophages. T cells in blood was used as a positive control.

### Pulse chase

Male and female heterozygous CSF1R^Mer-iCre-Mer^ tdTomato were fed tamoxifen diet for four weeks to label resident cells. Blood was collected at day 0, 3 weeks, 4 weeks, and 8 weeks after tamoxifen removal. Peripheral nerves from sciatic nerves, fascial nerves, vagal nerves, and DRG were pooled (PNS) and examined along with pooled brain and spinal cord (CNS) by flow cytometry at day 0 and 8 weeks following tamoxifen removal.

### RNAseq and data analyses

Library preparation, RNA-sequencing, data generation and quality-control was conducted by the ImmGen Consortium according to the consortium’s standard protocols (https://www.immgen.org/Protocols/ImmGenULI_RNAseq_methods.pdf). In short, the reads were aligned to the mouse genome GRCm38/mm10 primary assembly and gene annotation vM16 using STAR 2.5.4a. The raw counts were generated by using featureCounts (http://subread.sourceforge.net/). Normalization was performed using the DESeq2 package from Bioconductor. Differential gene expression analysis was performed using edgeR 3.20.9 in a pairwise manner among all conditions, and a total of 12,241 differentially expressed genes were defined with a p-value ≤ 0.001 and ≥ 4-fold difference. To construct the correlation plot, Euclidean distance among samples were calculated based on the differential expression matrix and clustering was performed using the ward.D2 algorithm in R. CNS/PNS shared, PNS-specific and CNS-specific genes were determined by subclustering the differentially expressed genes based on the expression pattern with a refined k-mean clustering using R and followed by manual curations. For neuronal microenvironment analysis, only the transcriptome profile of macrophages and microglia from PNS and CNS were used and analyzed through the same pipeline as mentioned above.

### Comparison of published microglia data

External datasets for circos plot were obtained from Krasemann et al. Genes that are enriched in SOD1G93A, aging brain, MFP2-/-, brain irradiation, Alzheimer’s disease (5XFAD) and phagocytic microglia conditions were previously generated in Krasemann et al. By comparing the PNS and CNS-enriched genes with the disease signatures, we were able to define the number of conditions shared by the genes and coined the term as “connectivity”. Only genes with connectivity of 2 or above are shown in the circos plot. For the comparison with Sall1−/− data set (24), the log_2_ fold change of genes between CNS microglia and PNS macrophages were calculated and compared against the public data set. Correlation coefficient and p-value were calculated by lineregress in Scipy using Python3.

### Embryonic labeling

Homozygous CX3CR1-GFP female mice were rotated daily with CSF1R^Mer-iCre-Mer^ tdTomato male mice and checked for plugs in the morning. Plug-positive females were administered 1.5 mg of 4-Hydroxytamoxifen (Sigma, Cat. # H6278) and 1 mg of progesterone (Sigma, Cat. # P0130) dissolved in corn oil by oral gavage 8 days following identification of plugs in order to pulse the embryos at embryonic 8.5 days. Following birth, pups were immediately sacrificed. Blood was collected for flow cytometric analysis and tissues were fixed in PFA/sucrose for imaging.

### In situ hybridization

RNA in situ hybridization (ISH) was performed using the ViewRNA Tissue Assay Core Kit (Invitrogen Cat. # 19931) according to the manufacturer’s instructions, with a probe set designed for IL-34 and Prx.

### Cell preparation, 10X Single cell library preparation, sequencing and analyses

A total of 4,500 Flt3-negative and 16,000 Flt3-positive cells were loaded to separate lanes of the 10X Chip for preparation of two single-cell libraries. The library preparation was performed according to the manufacturer’s instructions (Chromium Single-cell v2; 10X Genomics, USA). A total of 153M and 174M reads were sequenced for Flt3-negative and Flt3-positive libraries respectively using Illumina HiSeq2500. Reads were mapped by using the cellranger pipeline v2.1.1 onto the reference genome grcm38/mm10. We filtered cells for those with ≥50,000 mapped reads, leaving ~1k Flt3 negative and 4k Flt3 positive cells. Downstream analyses were performed by using the package Seurat2 in R.

## References

1. Keren-Shaul, H., et al. (2017). “A Unique Microglia Type Associated with Restricting Development of Alzheimer’s Disease”. Cell 169(7): 1276–1290 e1217.

2. Li, Q. and B. A. Barres (2018). “Microglia and macrophages in brain homeostasis and disease”. Nat Rev Immunol 18(4): 225–242.

3. Frost, J. L. and D. P. Schafer (2016). “Microglia: Architects of the Developing Nervous System.” Trends Cell Biol 26(8): 587–597.

4. Deczkowska, A., et al. (2018). “Disease-Associated Microglia: A Universal Immune Sensor of Neurodegeneration.” Cell 173(5): 1073–1081.

5. Butovsky, O., et al. (2014). “Identification of a unique TGF-beta-dependent molecular and functional signature in microglia.” Nat Neurosci 17(1): 131–143.

6. Krasemann, S., et al. (2017). “The TREM2-APOE Pathway Drives the Transcriptional Phenotype of Dysfunctional Microglia in Neurodegenerative Diseases.”. Immunity 47(3): 566–581 e569.

7. Kaucka, M. and I. Adameyko (2014). “Non-canonical functions of the peripheral nerve.”. Exp Cell Res 321(1): 17–24.

8. Chandran, V., et al. (2016). “A Systems-Level Analysis of the Peripheral Nerve Intrinsic Axonal Growth Program.” Neuron 89(5): 956–970.

9. Klein, D. and R. Martini (2016). “Myelin and macrophages in the PNS: An intimate relationship in trauma and disease.”. Brain Res 1641(Pt A): 130–138.

10. Shepherd, A. J., et al. (2018). “Macrophage angiotensin II type 2 receptor triggers neuropathic pain.” Proc Natl Acad Sci U S A 115(34): E8057–E8066.

11. Cattin, A. L., et al. (2015). “Macrophage-Induced Blood Vessels Guide Schwann Cell-Mediated Regeneration of Peripheral Nerves.” Cell 162(5): 1127–1139.

12. Goodrum, J. F. and D. L. Novicki (1988). “Macrophage-like cells from explant cultures of rat sciatic nerve produce apolipoprotein E.”. J Neurosci Res 20(4): 457–462.

13. Monaco, S., et al. (1992). “MHC-positive, ramified macrophages in the normal and injured rat peripheral nervous system.” J Neurocytol 21(9): 623–634.

14. Ginhoux, F., et al. (2010). “Fate mapping analysis reveals that adult microglia derive from primitive macrophages.” Science 330(6005): 841–845.

15. Wang, Y., et al. (2012). “IL-34 is a tissue-restricted ligand of CSF1R required for the development of Langerhans cells and microglia.” Nat Immunol 13(8): 753–760.

16. Pirzgalska, R. M., et al. (2017). “Sympathetic neuron-associated macrophages contribute to obesity by importing and metabolizing norepinephrine.” Nat Med 23(11): 1309–1318.

17. De Schepper, S., et al. (2018). “Self-Maintaining Gut Macrophages Are Essential for Intestinal Homeostasis.” Cell 175(2): 400–415 e413.

18. Gautier, E. L., et al. (2012). “Gene-expression profiles and transcriptional regulatory pathways that underlie the identity and diversity of mouse tissue macrophages.” Nat Immunol 13(11): 1118–1128.

19. Dick, S. A., et al. (2019). “Self-renewing resident cardiac macrophages limit adverse remodeling following myocardial infarction.” Nat Immunol 20(1): 29–39.

20. Di Liberto, G., et al. (2018). “Neurons under T Cell Attack Coordinate Phagocyte-Mediated Synaptic Stripping.”. Cell 175(2): 458–471 e419.

21. Lavin, Y., et al. (2014). “Tissue-resident macrophage enhancer landscapes are shaped by the local microenvironment.” Cell 159(6): 1312–1326.

22. Deming, Y., et al. (2019). “The MS4A gene cluster is a key modulator of soluble TREM2 and Alzheimer’s disease risk.” Sci Transl Med 11(505).

23. Friedman, B. A., et al. (2018). “Diverse Brain Myeloid Expression Profiles Reveal Distinct Microglial Activation States and Aspects of Alzheimer’s Disease Not Evident in Mouse Models.” Cell Rep 22(3): 832–847.

24. Buttgereit, A., et al. (2016). “Sall1 is a transcriptional regulator defining microglia identity and function.” Nat Immunol 17(12): 1397–1406.

25. Lund, H., et al. (2018). “Fatal demyelinating disease is induced by monocyte-derived macrophages in the absence of TGF-beta signaling.” Nat Immunol 19(5): 1–7.

26. Haimon, Z., et al. (2018). “Re-evaluating microglia expression profiles using RiboTag and cell isolation strategies.” Nat Immunol 19(6): 636–644.

27. Ziv, Y., et al. (2006). “Immune cells contribute to the maintenance of neurogenesis and spatial learning abilities in adulthood.” Nat Neurosci 9(2): 268–275.

28. Bennett, F. C., et al. (2018). “A Combination of Ontogeny and CNS Environment Establishes Microglial Identity.”. Neuron 98(6): 1170–1183 e1178.

29. Cronk, J. C., et al. (2018). “Peripherally derived macrophages can engraft the brain independent of irradiation and maintain an identity distinct from microglia.” J Exp Med 215(6): 1627–1647.

30. Gomez Perdiguero, E., et al. (2015). “Tissue-resident macrophages originate from yolk-sac-derived erythro-myeloid progenitors.” Nature 518(7540): 547–551.

31. Zhu, Y., et al. (2017). “Tissue-Resident Macrophages in Pancreatic Ductal Adenocarcinoma Originate from Embryonic Hematopoiesis and Promote Tumor Progression.” Immunity 47(3): 597.

32. Epelman, S., et al. (2014). “Embryonic and adult-derived resident cardiac macrophages are maintained through distinct mechanisms at steady state and during inflammation.” Immunity 40(1): 91–104.

33. Mueller, M., et al. (2003). “Macrophage response to peripheral nerve injury: the quantitative contribution of resident and hematogenous macrophages.” Lab Invest 83(2): 175–185.

34. Stratton, J. A., et al. (2018). “Macrophages Regulate Schwann Cell Maturation after Nerve Injury.” Cell Rep 24(10): 2561–2572 e2566.

35. Goldmann, T., et al. (2016). “Origin, fate and dynamics of macrophages at central nervous system interfaces.” Nat Immunol 17(7): 797–805.

36. Van Hove, H., et al. (2019). “A single-cell atlas of mouse brain macrophages reveals unique transcriptional identities shaped by ontogeny and tissue environment.” Nat Neurosci.

37. Li, Q., et al. (2019). “Developmental Heterogeneity of Microglia and Brain Myeloid Cells Revealed by Deep Single-Cell RNA Sequencing.” Neuron 101(2): 207–223 e210.

38. Kim, K. W., et al. (2016). “MHC II+ resident peritoneal and pleural macrophages rely on IRF4 for development from circulating monocytes.” J Exp Med 213(10): 1951–1959.

39. Qian, B. Z., et al. (2011). “CCL2 recruits inflammatory monocytes to facilitate breast-tumour metastasis.” Nature 475(7355): 222–225.

40. Feltri, M. L., et al. (1999). “P0-Cre transgenic mice for inactivation of adhesion molecules in Schwann cells.” Ann N Y Acad Sci 883: 116–123.

41. Takasato, M., et al. (2004). “Identification of kidney mesenchymal genes by a combination of microarray analysis and Sall1-GFP knockin mice.” Mech Dev 121(6): 547–557.

